# Kinetic proofreading decouples signal strength and range in paracrine gradient formation

**DOI:** 10.64898/2026.07.17.739274

**Authors:** Purushottam D. Dixit, Aviral Jain

## Abstract

Spatial gradients of signaling molecules pattern multicellular tissues with high precision. The canonical synthesis-diffusion-degradation (SDD) framework imposes a tradeoff on these gradients: ligand-receptor interactions that generate downstream signaling activity are also responsible for consuming the ligand. Correspondingly, at a fixed ligand synthesis rate, raising ligand-receptor affinity increases local signal strength at the expense of spatial range, and lowering it extends range at the expense of strength. Recent live-imaging measurements appear to violate this seemingly fundamental tradeoff, with low-affinity ligands of the epidermal growth factor receptor (EGFR) diffusing farther *and* driving spatially broader signaling activity compared to high-affinity ligands. Here we explain these observations with a model of multi-step ligand processing at the receptor, and show that the activity–range tradeoff is a consequence of receptor architecture rather than a physical necessity. When receptors process ligand through a multi-step phosphorylation cascade with kinetic-proofreading-like resetting, the states that generate activity decouple from those that consume ligand, and signaling activity and range increase together over a finite window of ligand residence time. This lets cells tune how far a signal travels independently of how strongly it acts through tuning signaling parameters. Realistic EGFR parameters place the low-affinity ligands in this window. Because multi-site phosphorylation and preferential degradation of the active receptor recur across multiple receptor families, kinetic proofreading may be a general strategy for controlling signaling range.

**Significance Statement:** Cells coordinate by releasing molecules that bind receptors on neighboring cells. For a fixed supply, how strongly a signal acts and how far it spreads are locked together: tight binding gives a strong response but the molecule is captured near its source, while weak binding spreads farther but signals feebly. Yet recent imaging of epidermal growth factor receptor ligands shows the opposite: weak binders activate a broader field of cells. We show this limit reflects how receptors read the signal, not physics. A receptor that processes a bound molecule through several steps, and can release it partway, separates the states that signal from those that destroy it. Cells, and engineers, can then set a signal’s reach independently of its strength.

## I. INTRODUCTION

Spatial gradient formation is central to paracrine signaling across a range of biological processes including angiogenesis^1^, wound healing^2^, and immune responses^3^, where ligands from a localized source drive responses across tens to hundreds of micrometers^4^. Paracrine gradient formation is usually modeled within the synthesis-diffusion-degradation (SDD) framework^5–10^, in which a localized source, ligand diffusion, and receptor-mediated endocytosis and degradation produce a profile that decays away from the source. Two scales characterize the resulting signal: the activity *a*(*r*_*c*_) at the source boundary, which sets the maximum local response, and the decay length *λ*, over which the profile falls off into the surrounding tissue^4,9–14^.

A variety of molecular mechanisms, from alternative transport routes to signal-dependent feedback shape both activity and range^15–29^. Nonetheless, the two scales cannot be chosen independently. Several tradeoffs are known to constrain gradient formation^9,10,30,31^. Here we formulate an activity–range tradeoff that is implicit in the SDD framework but, to our knowledge, has not been stated explicitly, and that ties signaling output directly to reach. In SDD, the ligand-bound receptor that generates signaling activity is also the state that consumes the ligand. Therefore, at a fixed source strength, increasing ligand-receptor affinity raises the local signaling activity albeit at the expense of the spatial range. Conversely, decreasing ligand affinity extends the range at the expense of the activity.

Recent live-cell imaging in epithelial monolayers appears to violate this tradeoff in epidermal growth factor receptor (EGFR) signaling. Using single producer cells surrounded by a field of receiver cells, Deguchi et al.^32^ found that low-affinity ligands epiregulin (EREG) and amphiregulin (AREG)^33^ drive ERK activation, a downstream readout of receptor activity, that is spatially broader compared to activation downstream of high-affinity ligands such as HBEGF. As the authors emphasized^32^, while low-affinity ligands are expected to reach farther^34,35^, they are also expected to activate EGFR more weakly, so the resulting activation profiles should be weaker as well. Yet a broader activation profile means that appreciable signaling activity, not merely the diffusing ligand, reaches farther from the source, so range and activity appear to increase together as affinity decreases, in tension with the SDD tradeoff.

How can activity and range both increase with decreasing ligand affinity? The activity–range tradeoff in SDD follows from treating the receptor as a passive sink, in which binding leads directly to activation and degradation. In contrast, real receptors are not passive: receptor tyrosine kinases, GPCRs, JAK-STAT receptors, and immunoreceptors undergo multi-site phosphorylation, kinetic proofreading, and other posttranslational modifications after ligand binding, and different receptor states contribute differently to downstream activity and to endocytosis and degradation^36–41^. Non-equilibrium receptor processing and spatial kinetic proofreading have been examined for noise-robust positional inference and for substrate discrimination^42–44^, but how they shape activity gradients has not been explored.

Here we show that ubiquitous receptor architectures can potentially break the activity–range tradeoff that is fundamental to SDD models. We model a receptor that processes bound ligand through a multi-step phosphorylation cascade in which the ligand can dissociate from any state, returning the receptor to its free form, while only the fully phosphorylated state is committed to enhanced degradation. This is the kinetic proofreading motif introduced by Hopfield^45^ and Ninio^46^ and familiar from ligand discrimination by the T-cell receptor^40,41,47^, with one difference: rather than withholding signaling until the cascade completes, the receptor signals from the first phosphorylation and uses completion to trigger its own removal^36,48^. Signaling activity is then spread across all phosphorylated states while ligand consumption is dominated by the terminal one, so the two decouple. We refer to this architecture as gradient amplification via receptor proofreading (GARP).

Decoupling activity from consumption creates a finite window of ligand residence times (ligand-receptor affinities) in which local activity and gradient range both increase with decreasing ligand residence time, inverting the tradeoff. The proofreading motif is essential for this phenomena; a multi-step receptor that cannot release ligand partway remains constrained by the activity–range tradeoff. The position and depth of the window are set by the kinase rate and by the number of phosphorylation sites rather than by ligand affinity, so a cell can tune how far a signal travels independently of how strongly it acts; we develop both into testable predictions.

With realistic EGFR parameters, the low-affinity ligands EREG and AREG lie in or near this window, accounting for the apparent violation of the activity–range tradeoff by EGFR ligands. Because multi-step phosphorylation and preferential degradation of the active receptor recur across multiple receptor families, kinetic proof-reading may be a general strategy for setting signaling range.

## II. RESULTS

### A. A reaction-diffusion model for paracrine gradient formation

To explore the relationship between activity and spatial range and its dependence on ligand affinity, we develop a two-dimensional reaction-diffusion model of ligand transport and receptor activation in a planar tissue. A circular source of radius *r*_*c*_ at the origin releases ligand at constant flux *j*_0_ (molecules per unit time), which diffuses in two dimensions with diffusion constant *D* and is consumed by cell-surface receptors (Fig. 1A); we consider steady state, where source production balances receptor-mediated capture and degradation. The two-dimensional geometry captures the planar setting of the EGFR-ligand reach measurements that motivate this work, as well as of many other paracrine systems^12,28,29,32^.

**FIG. 1.**
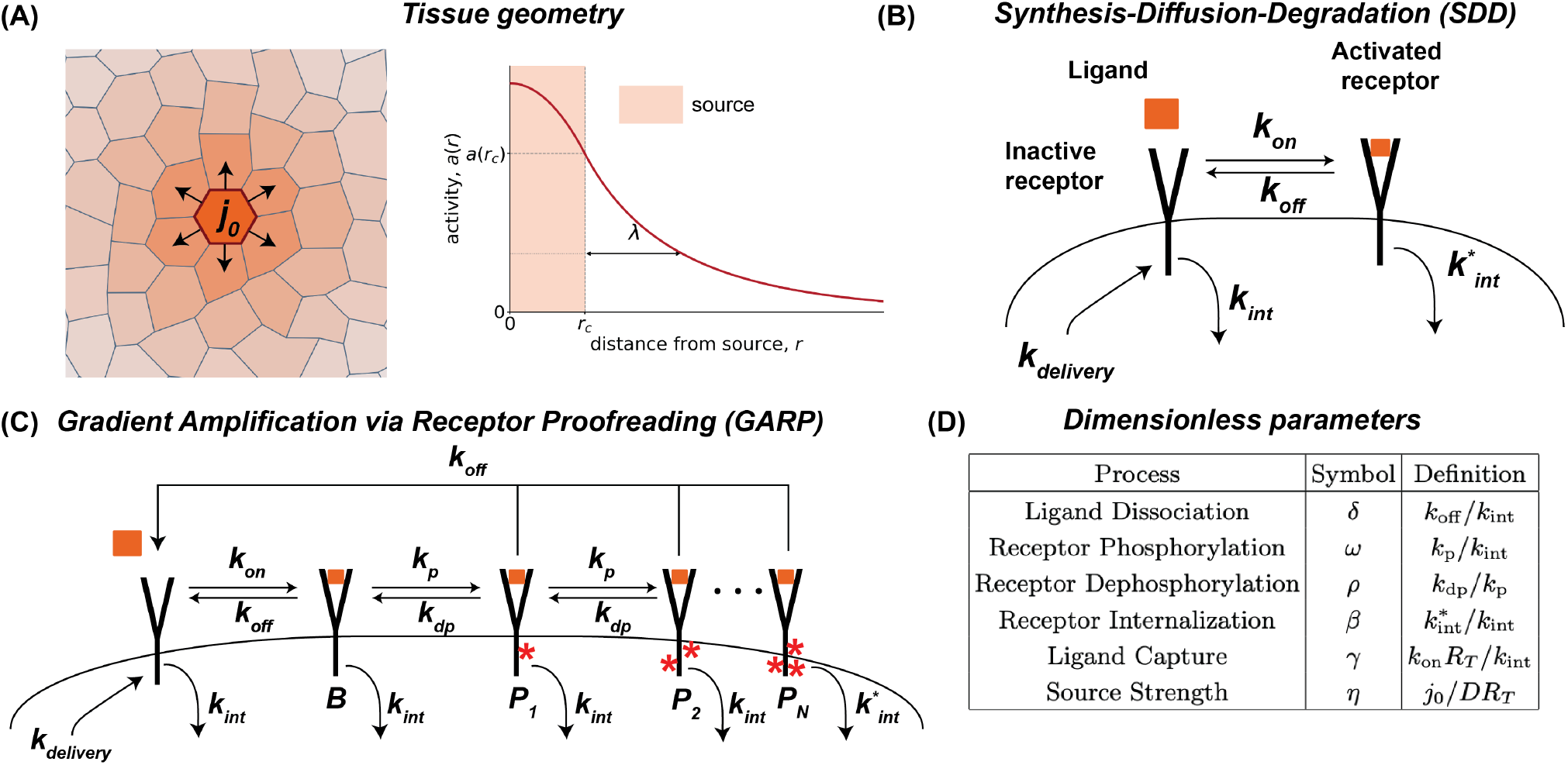
A reaction–diffusion model for paracrine gradient formation. (A) Tissue geometry. A circular source of radius *r*_*c*_ centered at the origin releases ligand at flux *j*_0_ into a planar tissue; ligand diffuses with coefficient *D* and is consumed by receptors. (B) SDD architecture: a single bound state both generates activity and is preferentially degraded. A free receptor *R* binds ligand at rate *k*_on_ and dissociates at rate *k*_off_ to form a single bound state *B*, which doubles as the active state and is internalized at the enhanced rate *β k*_int_. (C) GARP architecture (*N* -step phosphorylation cascade with proofreading). Ligand binding initiates a chain of phosphorylation steps *B* → *P*_1_ → · · · → *P*_*N*_ at rate *k*_*p*_; phosphorylated states can dephosphorylate (*P*_*n*_ →*P*_*n*−1_) at rate *ρ k*_*p*_; ligand can dissociate from any liganded state at rate *k*_off_ (the proofreading reset that gives GARP its name); only the terminal state *P*_*N*_ is preferentially internalized at the enhanced rate *β k*_int_. Signaling activity is the sum of all phosphorylated states. (D) Dimensionless parameters: *δ* = *k*_off_ */k*_int_, *ω* = *k*_*p*_*/k*_int_, *ρ* = *k*_dephos_*/k*_*p*_, 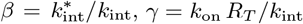, and *η* = *j*_0_*/*(*DR*_*T*_ ).

We compare two receptor architectures (Fig. 1B and C). In the traditional SDD model^5–8^, ligand binds a free receptor *R* to form a single bound state *B* that is both the active state and the state committed to enhanced internalization at rate *β k*_int_, where *k*_int_ is the basal internalization rate; free receptors are replenished to maintain steady state. The signaling activity at position 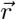 is the surface density of bound receptors, 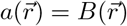.

In the gradient amplification via receptor proofreading (GARP) architecture (Fig. 1C), the ligand-bound receptor is sequentially phosphorylated at multiple sites, *B* →*P*_1_→ · · · →*P*_*N*_, at per-step rate *k*_*p*_, with reverse steps at rate *ρ k*_*p*_; here *B* is the unphosphorylated bound state and *P*_*n*_ the *n*-th phosphorylated state. Only the terminal state *P*_*N*_ is preferentially internalized, at *β k*_int_ with *β* ≥ 1; all other states are internalized at the basal rate. The defining feature of this architecture is the non-equilibrium proofreading motif^36,40,48^: from any liganded state, the ligand can dissociate at rate *k*_off_, returning the receptor to its free state *R* and the ligand to the diffusing pool. We define the signaling activity as the total surface density of phosphorylated receptors, 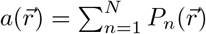.

The two architectures differ in one decisive respect. In SDD, a single state both generates activity and consumes ligand, so the two are inseparable. In GARP, activity is spread across all phosphorylated states, whereas ligand consumption is dominated by the terminal state *P*_*N*_ through its enhanced internalization. Activity and ligand loss are therefore carried by different combinations of receptor states, and as we show below, this is what allows them to be partially decoupled.

To compare ligands and architectures on a common footing, we nondimensionalize the equations, rescaling length by the diffusion-internalization length 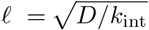 and ligand concentration by *c*_0_ = *j*_0_*/D* (SI Section I A). The receptor architecture is then set by the dissociation rate *δ* = *k*_off_ */k*_int_, the phosphorylation rate *ω* = *k*_*p*_*/k*_int_, the dephosphorylation ratio *ρ* = *k*_dephos_*/k*_*p*_, the preferential-internalization factor *β* ≥ 1, and the cascade depth *N*, collected as ***θ*** = (*δ, ω, ρ, β, N*); a capture rate *γ* = *k*_on_*R*_*T*_ */k*_int_ and a source strength *η* = *j*_0_*/*(*DR*_*T*_ ) complete the parameter set (Fig. 1D).

In a steady state with low ligand occupancy (SI Section I B; parameter ranges in SI Section I E), the dimensionless ligand profile satisfies a linear modified Helmholtz equation (SI Section I B),

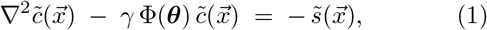

where 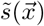 is the source distribution normalized to unit area integral, and Φ(***θ***) is a dimensionless *ligand loss factor* that encodes how the receptor architecture converts local ligand concentration into ligand consumption. Solving Eq. (1) for a uniform circular source of dimensionless radius 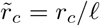 gives the closed-form concentration profile (SI Section I D)

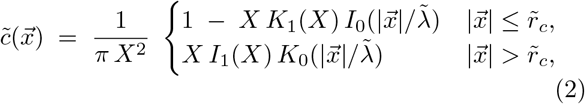

where 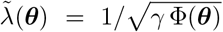 is the dimensionless decay length, *X* = *r*_*c*_*/λ* is the source radius measured in units of the decay length, and *I*_*n*_ and *K*_*n*_ are modified Bessel functions of the first and second kind.

In the low-occupancy regime, the signaling activity at every tissue position is proportional to the local ligand concentration

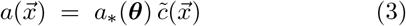

where *a*_∗_(***θ***) depends on the receptor processing architecture; the spatial structure of 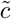 is otherwise universal.

The architecture-dependent quantities Φ(***θ***) and *a*_∗_(***θ***) take a simple form for the SDD model. Both Φ_SDD_(*δ, β*) and *a*_∗,SDD_(*δ, β*) can be written in closed form (SI Section I C),

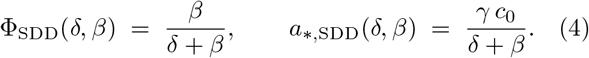

For GARP, Φ(***θ***) and *a*_∗_(***θ***) do not admit such compact closed forms; they follow from the coupled steady-state populations of the phosphorylated states and can be evaluated by a backward recurrence relationship (SI Section I C, Eqs. S32 and S33). Because activity (∑_*n*_ *P*_*n*_ ) and ligand loss 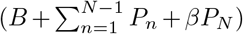 are carried by different combinations of receptor states, *a*_∗_ and Φ are no longer proportional, unlike in SDD.

### B. The activity–range tradeoff in SDD

We first illustrate the activity–range tradeoff in SDD (dashed lines in Fig. 2). The activity profile *a*(*r*)*/R*_*T*_ decreases approximately exponentially away from the source. Sweeping across a family of ligands with different dissociation rates *δ* shows that strong-binding ligands (low *δ*) produce sharp profiles with high source-boundary amplitude but a short reach, whereas weak-binding ligands (high *δ*) produce shallow profiles with low amplitude and long reach (Fig. 2A, dashed). The tradeoff is made quantitative by plotting the normalized local activity *a*(*r*_*c*_)*/R*_*T*_ against the normalized range *λ/r*_*c*_ as ligand affinity *δ* is swept (Fig. 2B, dashed), for several values of *β*. Each sweep traces a single monotonic curve: increasing *δ* trades activity for range, and varying the preferential-internalization factor *β* rescales the relationship without breaking it.

**FIG. 2.**
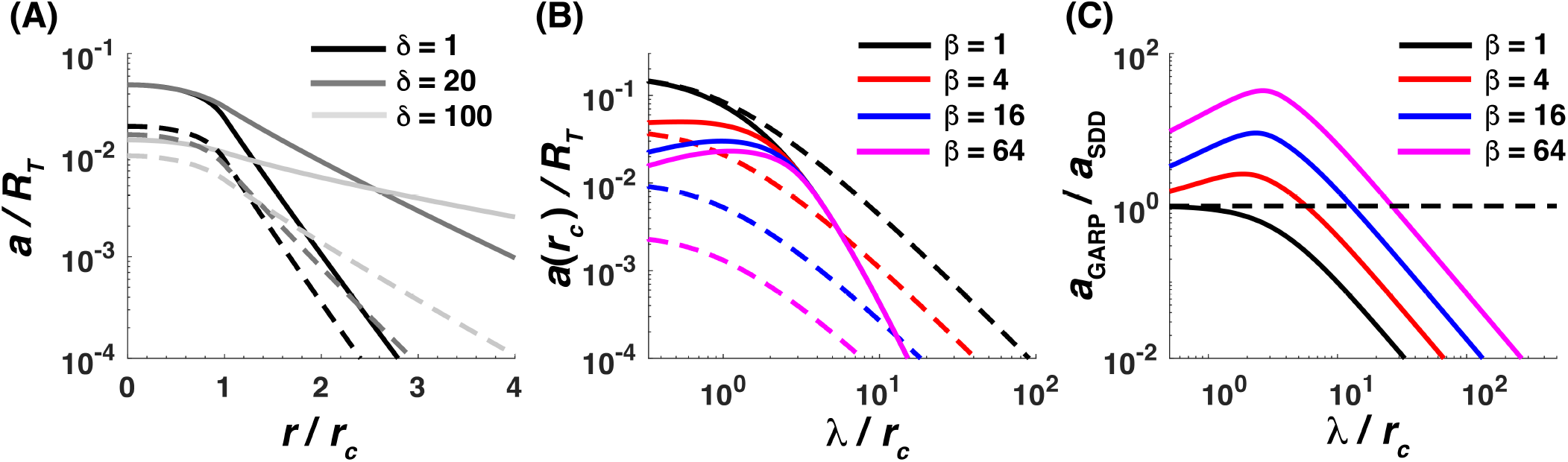
GARP overcomes the activity–range tradeoff. (A) Steady-state activity profiles *a*(*r*)*/R*_*T*_ versus scaled distance *r/r*_*c*_ at *β* = 16 for *δ* = 1, 20, 100, for SDD (dashed) and GARP (solid). In SDD, increasing *δ* (lowering affinity) lowers the overall activity profile while extending its spatial range. In contrast, in GARP, increasing *δ* into the amplification window extends the gradient with little loss of source-boundary activity. (B) Active receptor fraction at the source boundary, *a*(*r*_*c*_)*/R*_*T*_, versus the dimensionless gradient range *λ*(***θ***)*/r*_*c*_, parameterized by *δ*, for the SDD model (dashed lines) and the GARP architecture (solid lines) at *β* ∈ {1, 4, 16, 64} . (C) Amplification of local activity in GARP relative to SDD at matched range, *a*_GARP_*/a*_SDD_, versus *λ*(***θ***)*/r*_*c*_ for the same *β* values; the dashed line marks unity. Fixed parameters *γ* = 100, 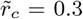, *η* = 0.1, *ω* = 100, *ρ* = 0.1, *N* = 10.

The tradeoff arises from a structural feature of the SDD architecture: the ligand-bound state *B* simultaneously carries the signaling activity (*a* = *B*) and is responsible for ligand loss (*β k*_int_ *B*). This dual role *couples* the activity scale to ligand loss flux: from Eq. (4), *a*_∗,SDD_ = (*γ c*_0_*/β*) Φ_SDD_, so *a*_∗_ and Φ share the same *δ* dependence up to constants. Combined with the scaling *λ*^2^ ∝ 1*/*Φ, this coupling between activity and ligand loss forces the source-boundary activity and the gradient range onto a single constraint surface.

### C. Activity and range increase together over a window of ligand affinity

The GARP architecture breaks this tradeoff that constrains gradient formation in SDD. As seen in Fig. 2A, raising the dissociation rate from *δ* = 1 to *δ* = 20 (lowering affinity) extends the range of the gradient while the source-boundary activity is essentially preserved, and only at very high *δ* = 100 does the boundary activity fall. In contrast, in SDD, the same gain in range is compensated by a proportional loss of activity (Fig. 2A, dashed). This is visible directly in the *a*(*r*_*c*_)–*λ/r*_*c*_ relationship (Fig. 2B): the GARP curves (solid) lift above the SDD curves (dashed) and develop an interior peak. As *δ* increases from strong binding, the curve first moves up and to the right, both source-boundary activity and range increasing together, and only past the peak does activity fall while range keeps growing. The height of the departure is set by *β*. At *β* = 1, activity and loss coincide again (except for the ligand-bound state which contributes to loss but not to activity) and GARP reduces to a tradeoff similar to the one observed in SDD. As *β* rises, activity *kinetically sorts itself* into the upstream phosphorylated states^48^ while the terminal state carries a growing share of ligand loss, so the peak lifts farther above the SDD curve. Panel C quantifies this as the ratio *a*_GARP_*/a*_SDD_ at matched range: for *β >* 1, the ratio exceeds unity across the short- and intermediate-range portion of the curve, reaching roughly 2.5-fold at *β* = 4, nearly tenfold at *β* = 16, and about 30-fold at *β* = 64 before crossing below unity at long range. At a fixed range, GARP thus delivers substantially more source activity than SDD, with the advantage growing in *β*.

With realistic parameter values, the observations by Deguchi et al. fall into place quantitatively. EGFR dissociation rates span *k*_off_ ∼ 10^−3^ s^−1^ for strong binders such as HBEGF and EGF to *k*_off_ ∼ 10^−1^ s^−1^ for EPGN, with EREG and AREG near the high-*k*_off_ end^48–51^. Taking *k*_int_ ∼ 10^−3^ s^−152,53^ and *k*_*p*_ ∼ 10^−1^ s^−149,51,54^ as representative EGFR-family values gives *δ*_HBEGF_ ∼ 1 and *δ*_EPGN_ ∼ 10 − 100 (SI Section I F), with *ω* ∼ 100 (SI Section I E). For these values the amplification optimum falls at *δ* ∼ 10 − 40, and therefore sits well above EGF and among EPGN, EREG, and AREG: EGF binds too tightly to reach the optimum, while the low-affinity ligands sit in or near the amplification window. This provides a quantitative account of the Deguchi et al.^32^ observation that low-affinity EGFR ligands produce broader ERK activation profiles than EGF. Since ERK activation is a downstream readout of receptor activity, this broader reach reflects a broader underlying receptor-activity profile, which is what GARP predicts.

### D. Kinetic proofreading is essential for gradient amplification

The activity–range tradeoff is not overcome by multi-step processing alone; the proofreading reset is required. Cascades that are structurally similar to GARP but lack the proofreading motif preserve the tradeoff. To confirm this, we compare the full GARP architecture against a variant in which dissociation occurs only from the ligand-bound state *B*; all phosphorylated receptors hold on to their ligands (SI Section I C 5). Figure 3 shows that without proofreading (dashed lines), the activity–range relationship is monotonic at every *β*, tracing the same kind of tradeoff curve as SDD. In contrast, full GARP (solid lines) develops the non-monotonic interior peak that breaks the tradeoff whenever *β >* 1; at *β* = 1 both architectures follow the activity–range tradeoff. The difference is best understood in the limit *ρ* →0, where the populations of adjacent states are related explicitly:

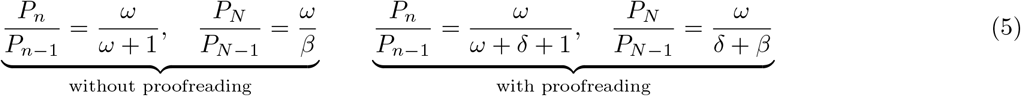

**FIG. 3.**
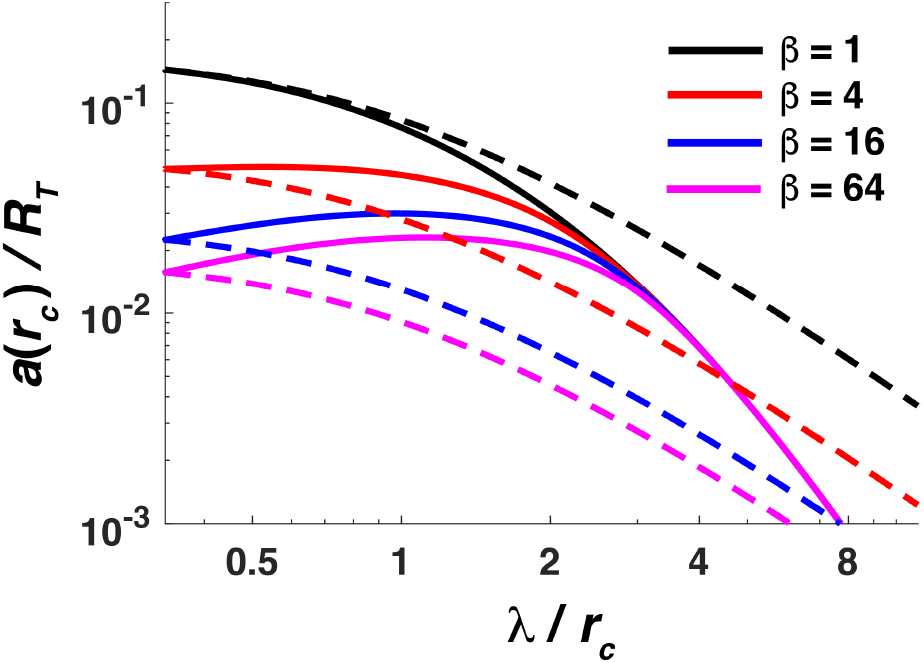
Kinetic proofreading is essential for gradient amplification. Active receptor fraction at the source boundary *a*(*r*_*c*_)*/R*_*T*_ versus dimensionless gradient range *λ*(***θ***)*/r*_*c*_, parameterized by *δ*, for the no-proofreading cascade (dashed) and full GARP (solid) at *β* ∈ {1, 4, 16, 64} . Full GARP develops a non-monotonic interior peak for *β >* 1, whereas the no-proofreading variant is monotonic at every *β*, restoring the activity–range tradeoff. Fixed parameters *γ* = 100, 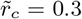, *η* = 0.1, *ω* = 100, *ρ* = 0.1, *N* = 10.

The contrast is immediate. Without proofreading, the ratios depend only on *ω* and *β*; the ligand affinity *δ* does not enter. The cascade occupancy is therefore fixed as *δ* is varied, *δ* scales only the overall bound-state amplitude, and the *a*(*r*_*c*_)–*λ/r*_*c*_ curve stays monotonic, as in SDD. With proofreading, every ratio picks up a *δ* term in the denominator, so the shape of the cascade occupancy changes with affinity. At large *δ* (weak binding, long range), upstream states empty out because most receptors dissociate before completing the cascade, so activity is low; at small *δ* (tight binding, short range), occupancy piles into the terminal state *P*_*N*_, which is preferentially internalized and consumes ligand quickly, again limiting activity. Activity is therefore maximized at intermediate *δ*, and this optimum is exactly the interior peak of the GARP curve in Fig. 3 that the no-proofreading cascade lacks. Proofreading, not the multi-step cascade alone, is what breaks the activity–range tradeoff.

### E. Signaling range is tunable through kinase activity and cascade length

The departure of GARP from SDD in the relationship between *a*(*r*_*c*_) and *λ*(***θ***)*/r*_*c*_, established above, is set by the receptor architecture itself. Two parameters control where the amplification window sits and how much amplification it delivers: the phosphorylation rate *ω* and the processing-chain length *N*. Because neither is a property of the ligand, they provide knobs by which a cell can set signaling range independently of ligand affinity.

The phosphorylation rate *ω* controls how far the amplification extends along the range axis, and at what cost in activity. Faster phosphorylation lets a receptor take several phosphorylation steps before the ligand dissociates, so upstream activity is still built up even for weakly bound, short-residence ligands. Increasing *ω* therefore extends the range over which activity is maintained: the *a*(*r*_*c*_)–*λ/r*_*c*_ curve flattens and its interior peak moves outward, so the gradient can reach farther while the source-boundary activity stays essentially unchanged. This reach is not free. The peak now sits at weaker binding (larger *δ*, and hence longer *λ*, since range grows monotonically with *δ*), where fewer receptors are ligand-engaged and the source-boundary amplitude is intrinsically lower, so the overall activity level of the curve drops as *ω* rises. In short, faster phosphorylation trades peak activity for range, the optimum moving to weaker binding as *ω* grows. Figure 4A illustrates this: as *ω* increases across {4, 16, 64, 256} (fixed *β* = 16, *N* = 10), an interior peak emerges once phosphorylation is fast enough to compete with dissociation, and thereafter shifts to longer range and lower activity.

**FIG. 4.**
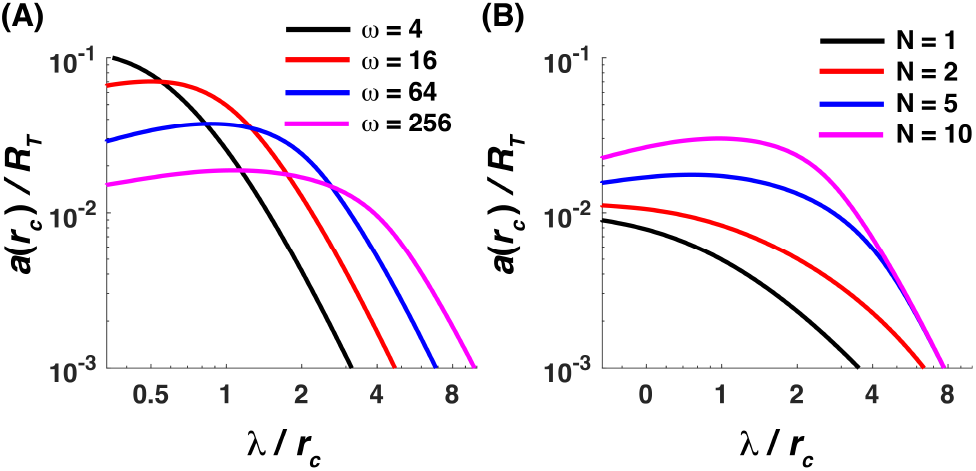
The amplification window is shaped by phosphorylation rate and chain length. Active receptor fraction at the source boundary *a*(*r*_*c*_)*/R*_*T*_ versus dimensionless gradient range *λ*(***θ***)*/r*_*c*_, parameterized by *δ* (solid lines). (A) Varying the phosphorylation rate *ω* ∈ {4, 16, 64, 256} at fixed *β* = 16, *N* = 10: an amplification peak emerges for larger *ω* and thereafter shifts toward larger *δ* and longer range, while the peak activity falls. (B) Varying the chain length *N* ∈ {1, 2, 5, 10} at fixed *ω* = 100, *β* = 16: the window opens only for larger *N*, and both the peak amplitude and its width grow with *N* . Other parameters *γ* = 100, 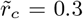, *η* = 0.1, *ρ* = 0.1.

The processing-chain length *N* sets how much amplification the architecture delivers and over how wide a range. Each additional phosphorylation step adds an upstream state that contributes to signaling activity but is internalized only at the basal rate, raising activity without adding to ligand loss. Lengthening the cascade therefore lifts the whole amplification peak: the *a*(*r*_*c*_)–*λ/r*_*c*_ curve rises to higher activity at its optimum, and because more intermediate states can accumulate occupancy, the peak also broadens, sustaining amplification over a wider range of *λ*. Figure 4B illustrates this: at fixed *ω* = 100, *β* = 16, sweeping *δ* for *N* ∈ {1, 2, 5, 10} shows the window opening only for larger *N*, with both the peak activity and the range-width of the window growing as *N* increases.

Both knobs yield experimental predictions. First, reducing the per-step phosphorylation rate *ω*, for example by mild kinase inhibition, should shift the optimum toward tighter-binding, longer-lived ligands (smaller *δ*, shorter *λ*) rather than uniformly suppressing signaling (Fig. 4A). Second, receptors engineered with altered numbers of phosphorylation sites, through truncated C-terminal tails or mutated tyrosines, should change both the magnitude and the position of the amplification window (Fig. 4B). Both predictions distinguish multi-step processing from a passive multi-state sink and from SDD, in which the ordering of ligand activities at the source boundary depends on *δ* but not on *ω* or *N*.

## III. DISCUSSION

We showed that a receptor which processes bound ligand through a multi-step phosphorylation cascade decouples source-boundary activity from gradient range, overcoming the activity–range tradeoff of the SDD framework. Because activity is the uniformly weighted sum over all phosphorylated states while ligand loss weights the terminal state by *β* ≫ 1, activity accumulates upstream without feeding the loss that pins the source flux, opening a finite residence-time window in which activity and range grow together. The proofreading reset, dissociation from any liganded state, is essential to this escape.

The mechanism composes with other gradient-shaping processes rather than replacing them. Self-enhanced degradation^23^, transcytosis^55^, and signaling-driven autocrine relays that generate ERK activation^28,29^ act on ligand transport or on receptor density, and would combine with the receptor-side accumulation described here.

The architecture also has consequences for noise. Because sequential phosphorylation accumulates evidence of ligand binding while terminal-state degradation both reports and consumes it, the receptor acts as an integral-feedback low-pass filter^51,56^, with an averaging time ∼ *N/k*_*p*_ set by the time to traverse the cascade. Rapid fluctuations in extracellular ligand are filtered out while sustained binding accumulates, buffering signaling near sparse or episodic sources; a full stochastic treatment remains for future work.

Several limitations are worth noting. We treat only the steady-state regime, whereas the Deguchi ERK measurements are temporally propagating waves^32^; their activated area is thus a thresholded, dynamic proxy for our graded steady-state activity profile, and a full account would require a time-resolved treatment. We coarse-grain EGFR-specific complications, including ErbB heterodimerization and the sorting of internalized receptors between recycling and degradation^33,50^; the ligand-dependent sorting reported by Deguchi et al.^32^ is itself a residence-time effect, which our model absorbs into the internalization rates *k*_int_ and *β k*_int_. We hold the receptor density *R*_*T*_ constant, neglecting signaling-dependent receptor turnover. We isolate the receptor-level contribution to ERK activity and omit cell–cell contact inputs. Finally, we treat the tissue as a continuum and neglect ligand-concentration noise, which is small for ligand retained on the basal surface of a confluent monolayer and unlikely to differ between ligands in a way that could explain the reach differences.

More broadly, the mechanism requires only three ingredients: multi-step receptor modification following ligand binding, a per-step modification rate comparable to or faster than the dissociation rate, and preferential internalization of the terminal state. These conditions are met across receptor tyrosine kinases, GPCRs, JAK-STAT receptors, and immunoreceptors^36–41^. Wherever they hold, the reach of a paracrine signal is set not by the affinity of the ligand but by how the receptor processes it: a cell can tune how far a signal travels independently of how strongly it acts, through kinase activity and receptor architecture rather than by changing the ligand itself. Kinetic proofreading may therefore be a general strategy for controlling how far signals travel.

## I. SUPPLEMENTARY MATERIALS

### A. Derivation of the reaction-diffusion equation

We write the dimensional rate equations for both receptor architectures (SDD and the *N*-step phosphorylation cascade), derive the receptor conservation relations, rescale to dimensionless form, and show that the low-occupancy steady state reduces the receptor-coupled system to the linear modified Helmholtz equation used throughout the main text.

#### 1. Dimensional rate equations

The ligand concentration 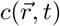 (molecules per unit tissue area) and the free-receptor surface density 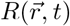 satisfy

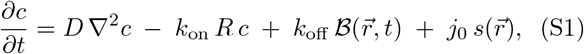

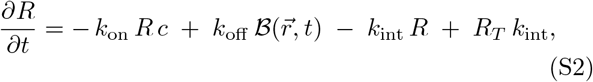

where *D* is the ligand diffusion coefficient, *k*_on_ and *k*_off_ the binding and dissociation rate constants, *k*_int_ the basal receptor internalization rate, *j*_0_ the total source flux (molecules per unit time), 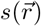 the normalized source distribution (∫*s d*^2^*r* = 1), *R*_*T*_ the resting total receptor surface density, and 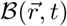 the total concentration of species that can dissociate the ligand. The trafficking flux *R*_*T*_ *k*_int_ guarantees *R* → *R*_*T*_ in the absence of ligand.

For the SDD model, ℬ = *B* (a single liganded state), and the bound state satisfies

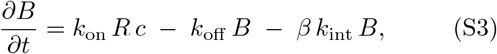

with *β k*_int_ the enhanced internalization rate of the active state.

For the *N*-step phosphorylation cascade, 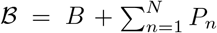, and the bound and phosphorylated states satisfy

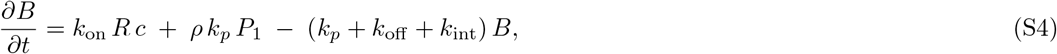

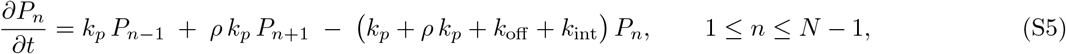

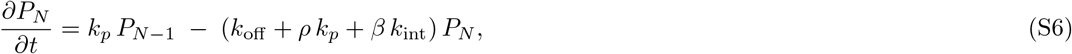

with the convention *P*_0_ ≡ *B* in the recursion. Phosphorylation occurs at rate *k*_*p*_, dephosphorylation at rate *ρ k*_*p*_, dissociation from any liganded state at rate *k*_off_, basal internalization of intermediate states at *k*_int_, and preferential internalization of the terminal state at *β k*_int_ (*β* ≥ 1).

#### 2. Receptor conservation relation

Summing all receptor equations and cancelling the binding/dissociation, phosphorylation, and dephosphorylation pairs gives

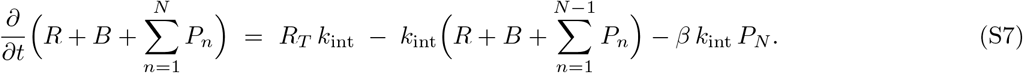

At steady state,

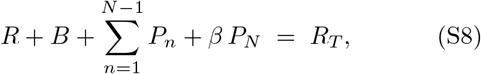

which reduces to *R* +*β B* = *R*_*T*_ for the SDD model. The weight on each state in Eq. (S8) is exactly the rate at which the state is internalized, divided by the basal rate.

#### 3. Dimensionless rescaling

To simplify the model representation, we identify several dimensionless quantities. We rescale time by the basal internalization rate, *τ* = *k*_int_ *t*; length by the diffusion-internalization length 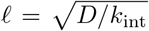, with 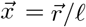 ligand concentration by the source-driven scale *c*_0_ = *j*_0_*/D*, with 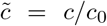 and receptor surface densities by *R*_*T*_, with *ψ* = *R/R*_*T*_, *b* = *B/R*_*T*_, *p*_*n*_ = *P*_*n*_*/R*_*T*_ . The source distribution rescales as 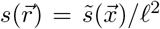 with 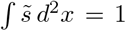. This rescaling produces the dimensionless groups

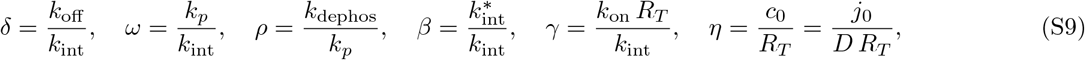

together with the cascade depth *N*. The receptor parameters are collected into ***θ*** = (*δ, ω, ρ, β, N*). The dimensionless equations are

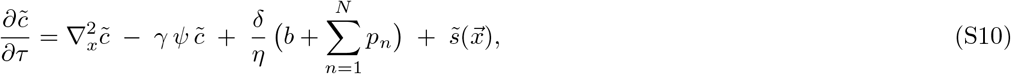

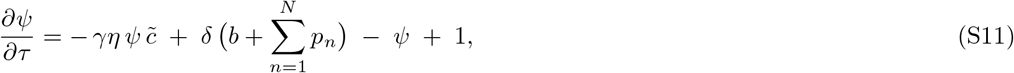

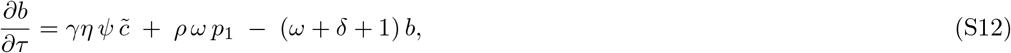

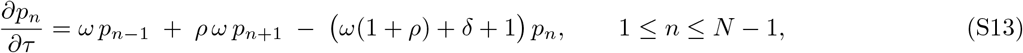

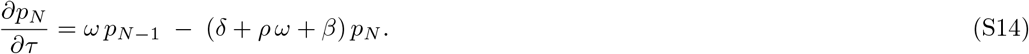

#### 4. Steady-state reduction of the ligand PDE

Dividing the summed liganded-state equations by *η* and adding the ligand equation, then cancelling the binding, dissociation, phosphorylation, and dephosphorylation terms, gives at steady state,

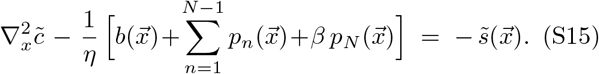

The only net ligand sinks are the internalization pathways, weighted by their respective rates. The binding/dissociation and phosphorylation/dephosphorylation steps shuttle receptors between states without removing ligand. By the receptor conservation relation (Eq. S8) in dimensionless form, the bracket is exactly 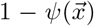, so that

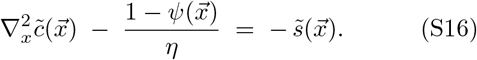

### B. Low-occupancy approximation

We work in the low-occupancy regime, defined by 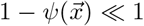 at every tissue position, where *ψ* = *R/R*_*T*_ is the free-receptor density measured against its resting value. By the steady-state balance (Eq. S8), 1 − *ψ* is the liganded-receptor fraction weighted by internalization rate, the terminal state carrying weight *β*; since *β* ≥ 1, it bounds the fraction of receptors engaged by ligand from above. Saturation is controlled by the dimensionless source strength *η* = *j*_0_*/*(*DR*_*T*_ ) (SI Section I E): the occupancy is 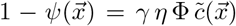, which tracks 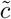 and is therefore largest at the center of the source. For representative EGFR values (*γ* ∼ 10^2^, *η* ∼ 0.1) it reaches ∼ 0.3 there, falls to ∼ 0.15 at the source boundary, and decays outward through the tissue, so the linearization remains controlled everywhere.

In this regime, we can replace 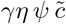 with 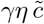 in the binding term of the receptor equations, since *ψ* ≈ 1 to leading order. The receptor steady-state equations then form a linear inhomogeneous system in 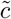:

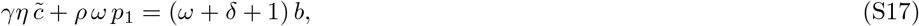

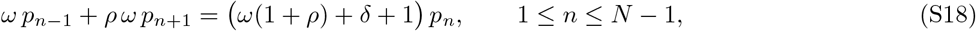

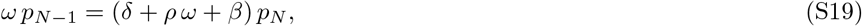

driven entirely by the inhomogeneous term 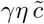 in the bound equation. The solutions are proportional to 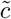:

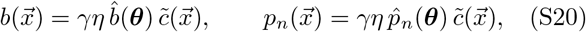

with 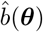 and 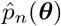 obtained by solving the homogeneous-recursion system with 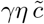 replaced by 1 in the bound equation. Closed-form expressions are derived in SI Section I C.

By the conservation relation,

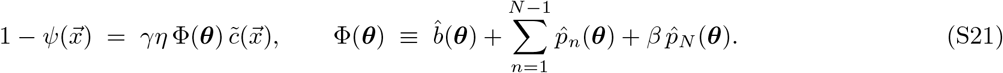

Substituting into the steady-state ligand PDE, *η* cancels, and the equation reduces to the linear modified Helmholtz form quoted in the main text:

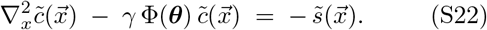

#### 1. Activity factorization

The dimensional signaling activity is

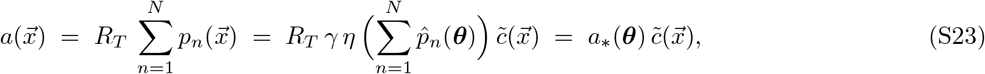

with

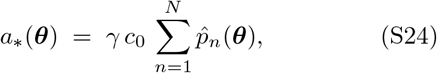

where we used *R*_*T*_ *η* = *c*_0_. The source flux *j*_0_ enters the activity scale linearly through *c*_0_ = *j*_0_*/D*, while the spatial structure of 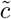 depends only on *γ*, Φ(***θ***), and the source geometry.

### C. Architecture-dependent quantities Φ(***θ***) **and** *a*_∗_(***θ***)

This section derives the explicit forms of the ligand loss factor Φ(***θ***) and the activity scale *a*_∗_(***θ***) that appear in the main-text Eqs. (1) and (3). Both quantities take simple closed forms for the SDD model and admit a compact downward recursion for the *N*-step phosphorylation cascade.

The amplitudes 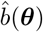 and 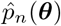 introduced in SI Section I A (Eq. S20) determine both Φ(***θ***) (Eq. S21) and *a*_∗_(***θ***) (Eq. S24). We now derive them in closed form.

#### 1. SDD limit: closed forms

In the SDD model, there are no phosphorylated states and *B* is internalized directly at the enhanced rate *β k*_int_. The bound-state equation becomes 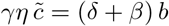, giving

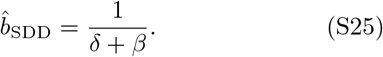

Activity in this architecture is 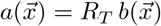, and from 1 − *ψ* = *β b*,

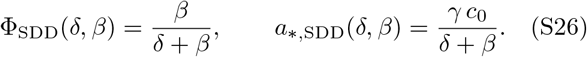

Φ_SDD_ is the fraction of bound-state residency that ends in internalization rather than dissociation.

#### 2. Cascade: downward recursion for 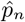

For the cascade we solve (S17)–(S19) by a downward recursion. Define the upstream-to-downstream state ratios

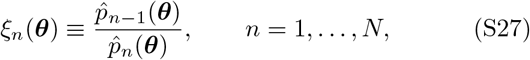

with 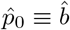. The terminal equation (S19) gives the starting value

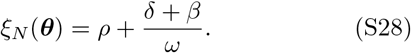

Dividing (S18) by 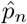 and using 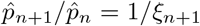 gives the downward recursion

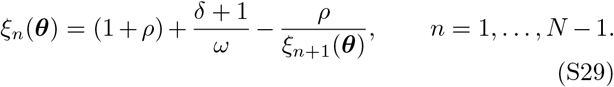

The bound-state equation (S17), normalized so that the right-hand side equals 1 and using 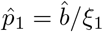, gives

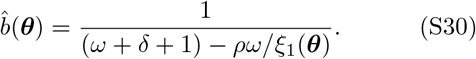

Defining the cumulative product 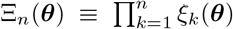 (with Ξ_0_ ≡ 1), the upstream amplitudes are

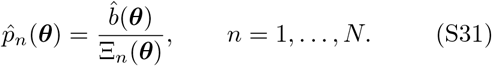

The recursion is solved by first setting *ξ*_*N*_ from (S28), then sweeping down through (S29) to obtain 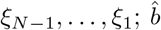 then follows from (S30) and the 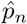 from (S31).

#### 3. Closed expressions for Φ(***θ***) **and** *a*_∗_(***θ***)

Substituting (S31) into the definitions (S21) and (S24) gives

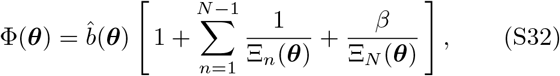

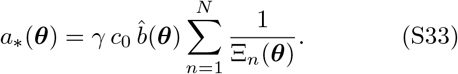

Both quantities depend on ***θ*** = (*δ, ω, ρ, β, N*) alone; the capture rate *γ* and source-flux scale *c*_0_ enter *a*_∗_ as overall prefactors.

#### 4. Limit checks

*a. Vanishing dephosphorylation (ρ* = 0). The recursion (S29) decouples: *ξ*_*n*_ = (*ω* + *δ* + 1)*/ω* for *n* = 1, …, *N* 1, and *ξ*_*N*_ = (*δ*+*β*)*/ω*. Then 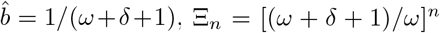 for *n < N*, and Ξ_*N*_ = [(*ω* + *δ* + 1)*/ω*]^*N*−1^ (*δ* + *β*)*/ω*.

*b. No preferential internalization (β* = 1). Summing (S17)–(S19) gives 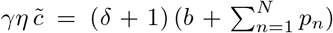, so 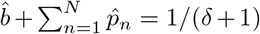. Substituting into (S21) with *β* = 1,

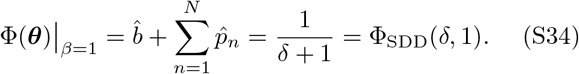

The cascade and SDD share the same ligand loss factor at *β* = 1, so the gradient ranges coincide. The activity scales differ by the unphosphorylated contribution, 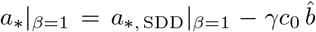, with the offset 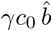 vanishing as *ω* → ∞ for *ρ* ≪ 1 (where the cascade clears *B* rapidly into the phosphorylated states and does not return it).

*c. No cascade (N* = 0). With no phosphorylated states, *B* is the only liganded state and is internalized at the enhanced rate *β k*_int_. The bound-state equation reduces to 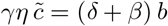, recovering (S26) with 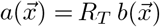 identified as the activity.

#### 5. Removing the proofreading reset

The no-proofreading cascade used in Fig. 3 is obtained by permitting dissociation only from the unphosphorylated bound state *B*: once a receptor is phosphorylated it retains its ligand until internalization. Only the phosphorylated-state equations change, losing their *δ* terms; the bound-state equation (S17) is untouched, since *B* still dissociates,

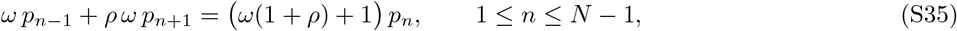

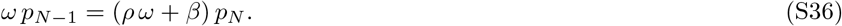

The downward recursion carries through with *δ* deleted from (S28) and (S29),

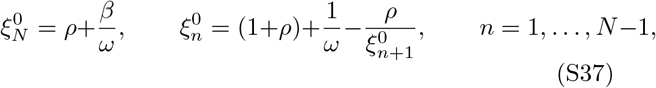

while 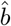 retains its *δ* dependence through (S30) with 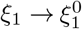.

The consequence is structural. The ratios 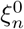, and hence the cumulative products 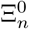, carry no *δ*: ligand affinity no longer reaches the cascade, and the shape of the phosphorylated-state occupancy is frozen as *δ* is varied. Equations (S32) and (S33) then reduce to 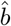 multiplied by *δ*-independent factors, and 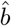 cancels between them,

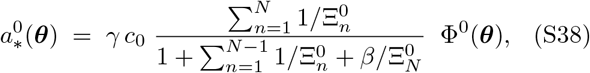

with the prefactor a function of (*ω, ρ, β, N*) alone. The activity scale is therefore strictly proportional to the ligand loss factor, exactly as in SDD, where *a*_∗,SDD_ = (*γc*_0_*/β*) Φ_SDD_ from (S26). Varying ligand affinity moves the system along this proportionality rather than off it, which is why the no-proofreading cascade traces a monotonic activity–range curve at every *β* (Fig. 3). The relation (S38) holds for any *ρ*; the main-text state ratios (Eq. 5), quoted in the transparent limit *ρ* → 0, are its special case.

### D. Bessel solution of the modified Helmholtz equation on a disk

This section derives the closed-form Bessel expression for the dimensionless ligand profile 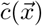 that appears in the main-text Eq. (2). The starting point is the modified Helmholtz equation

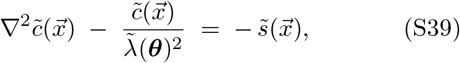

where 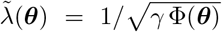 is the dimensionless decay length and the source is a top-hat on the disk of dimensionless radius 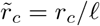, normalized to unit integral,

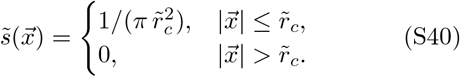

The problem is axisymmetric. We write 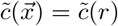 with 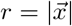.

#### 1. Inside and outside solutions

Inside the source 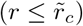, Eq. (S39) becomes

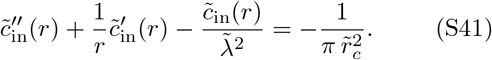

A particular solution is the constant 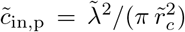. The homogeneous solutions of the modified Bessel equation in two dimensions are 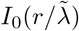 and 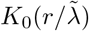 only *I*_0_ is regular at *r* = 0. Hence

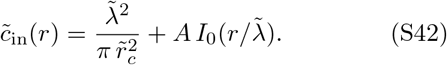

Outside the source 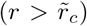, the equation is homogeneous, and only *K*_0_ decays at infinity:

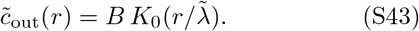

#### 2. Matching and the Wronskian identity

The source distribution is bounded (no delta function), so 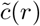 and its first derivative are continuous at 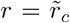 Using 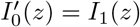 and 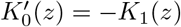, the derivative-matching condition is

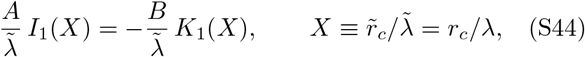

giving *B* = − *A I*_1_(*X*)*/K*_1_(*X*). The value-matching condition is

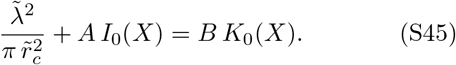

Substituting *B* and rearranging,

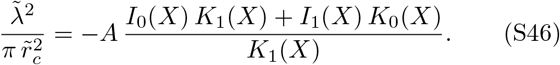

The combination in the numerator is the Wronskian of the modified Bessel functions,

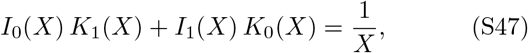

which gives 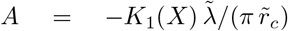 and 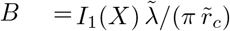.

Substituting back into (S42) and (S43) and using 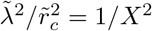,

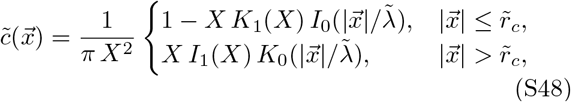

recovering the main-text Eq. (2).

#### 3. Boundary value and asymptotic limits

*a. Source-boundary value*. Evaluating (S48) at 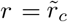 using the Wronskian (S47),

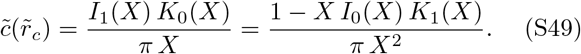

The signaling activity at the source boundary follows: 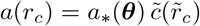

### E. Estimation of the dimensionless parameters

Beyond the dissociation rate *δ*, the model is set by the dimensionless groups *γ, β, ω, ρ*, and *η*, each a ratio of a receptor or transport rate to the basal internalization rate *k*_int_ ∼ 10^−3^ s^−152,53^. We estimate their order of magnitude for EGFR signaling in an epithelial monolayer; the values fix the operating regime rather than precise numbers.

The capture rate *γ* = *k*_on_*R*_*T*_ */k*_int_ uses the two-dimensional association constant appropriate to the planar geometry. Measured on-rates are three-dimensional (units of M^−1^s^−1^), whereas here ligand is confined to a thin pericellular layer of thickness *h*, so the effective areal constant is 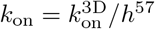 and

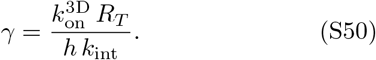

With an EGF–EGFR association rate 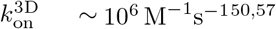, a receptor surface density *R*_*T*_ ∼ 10^2^– 10^3^ *µ*m^−257^, a pericellular thickness *h* ∼ 0.1–1 *µ*m, and *k*_int_ ∼ 10^−3^ s^−152,53^, we find *γ* ≫ 1: receptors capture ligand much faster than they internalize it. We use *γ* = 100 as a representative value.

The remaining groups are ratios of rates and need no geometric conversion. The preferential-internalization factor 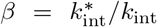 is the fold increase in internalization of the fully active receptor over the resting receptor; ligand-activated EGFR is internalized and degraded much faster than resting receptor, giving *β* of order 10^33,57^, and we sweep *β* ∈ {1, 4, 16, 64} to span this range. The phosphorylation rate *ω* = *k*_*p*_*/k*_int_ compares the per-step phosphorylation rate (*k*_*p*_ ∼ 10^−1^ s^−1^) to internalization, giving *ω* ∼ 10^2^. The dephosphorylation ratio *ρ* = *k*_dephos_*/k*_*p*_ is small when phosphorylation outpaces phosphatase turnover, of order 0–0.1; we use *ρ* = 0.1. The source strength *η* = *j*_0_*/*(*DR*_*T*_ ) sets the overall activity scale: for a small producing cluster shedding 10^2^–10^3^ ligands per second, *D* ∼ 10 *µ*m^2^s^−1^, and *R*_*T*_ as above, *η* ∼ 10^−2^–10^−1^, and we use *η* = 0.1. Together with *γ, η* also controls receptor saturation: the source-boundary occupancy 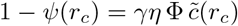 stays at or below ∼ 0.15 for these values, consistent with the low-occupancy approximation (SI Section I B).

These are order-of-magnitude estimates; they set the regime rather than precise values, and the central results, a finite amplification window that requires *β >* 1 and an optimum at intermediate *δ*, hold across the ranges above.

### F. EGFR ligand affinities

The EGFR ligand family spans more than two orders of magnitude in binding affinity, and the dimensionless dissociation rate *δ* = *k*_off_ */k*_int_ inherits this spread. Taking representative EGFR-family values for the basal internalization rate, *k*_int_ ∼ 10^−3^ s^−152,53^, together with measured off-rates^50^, strong-binding ligands such as EGF and HBEGF (*k*_off_ ∼ 10^−3^ s^−1^) map to small *δ*, of order unity, whereas weak-binding ligands such as EREG, AREG, and EPGN (*k*_off_ ∼ 10^−2^–10^−1^ s^−1^) map to large *δ*, of order 10–100. Because the amplification optimum falls at intermediate *δ* ∼ 10–40 for representative parameters, strong-binding ligands fall below the optimum while weak-binding ligands lie in or near the amplification window. These assignments are order-of-magnitude only: measured off-rates vary with cell type and assay, and ligand-specific trafficking, such as the endosomal dissociation of TGF-*α*, can alter a ligand’s effective residence time. We therefore use the mapping only to separate strong-from weak-binding ligands, not to place each ligand at a precise *δ*.

